# CVN636 potentiates orthosteric agonist responses selectively at mGlu_7/7_ homodimers

**DOI:** 10.64898/2026.10.17.759826

**Authors:** Xin Lin, Colleen M. Niswender, Wesley B. Asher, Jonathan A. Javitch

## Abstract

The metabotropic glutamate receptor 7 (mGlu_7_) is an emerging therapeutic target for central nervous system disorders, but progress has been limited by a lack of selective pharmacological tools. CVN636 was recently reported as a potent and selective allosteric agonist of this receptor, capable of receptor activation in the absence of glutamate, with demonstrated efficacy in a rodent model of alcohol use disorder. Here, using complementary bioluminescence resonance energy transfer-based assays that measure both downstream signaling and G protein recruitment, we show that CVN636 does not activate the receptor on its own. Instead, CVN636 acts as a positive allosteric modulator that enhances the potency and efficacy of orthosteric agonist responses, including those of glutamate, in an agonist-dependent manner. Blockade of orthosteric agonist binding with a competitive antagonist abolished CVN636 activity, consistent with this mechanism. Furthermore, using an assay that distinguishes signaling from defined receptor dimers, we find that CVN636 activity is selective for mGlu_7/7_ homodimers and does not extend to any mGlu_7_-containing heterodimer tested. Thus, in our assay system CVN636 is an mGlu_7/7_ homodimer-selective positive allosteric modulator of this receptor, suggesting that its i*n vivo* actions likely stem from action at these homomers.

## INTRODUCTION

Among the eight metabotropic glutamate (mGlu) receptors, mGlu_7_ is both the most widely expressed and the most evolutionarily conserved subtype in the mammalian central nervous system (CNS).^1^ Localized predominantly at presynaptic active zones of glutamatergic and GABAergic terminals, mGlu_7_ couples to Gα_i/o_ proteins to suppress neurotransmitter release and thereby modulate synaptic transmission across diverse brain circuits.^2–4^ A distinguishing feature of mGlu_7_ is its very low affinity for glutamate, requiring high micromolar to millimolar concentrations for activation — orders of magnitude higher than other mGlu subtypes. This has led to the suggestion that mGlu_7_ acts as a high-threshold sensor of glutamate overflow.^3,5,6^

Genetic and preclinical studies have implicated mGlu_7_ dysfunction in a range of CNS disorders, including autism spectrum disorder, attention-deficit hyperactivity disorder, epilepsy, depression, schizophrenia, and addiction.^7–15^ However, progress in fully delineating the therapeutic potential of mGlu_7_ has been hampered by a lack of potent and selective pharmacological tools.^16,17^ Existing activators, including allosteric agonists and several positive allosteric modulators (PAMs), have been limited by off-target activities, receptor internalization, insufficient CNS penetrance, or lack of subtype selectivity, complicating their use both as tool compounds and as therapeutic leads.^18–20^

In this context, the recent discovery of the allosteric mGlu_7_ compound CVN636 represented a significant advance.^21^ CVN636 was reported to be a highly potent (EC_50_ = 7 nM) allosteric agonist with selectivity for mGlu_7_ over the other mGlu receptor subtypes, favorable CNS penetrance, and no detectable off-target activity across 125 CNS-relevant targets. Importantly, acute oral administration of CVN636 reduced ethanol self-administration in a rodent model of alcohol use disorder, providing the first evidence of *in vivo* efficacy for a selective mGlu_7_ activator in an addiction paradigm. These properties positioned CVN636 as a first-in-class tool compound and potential drug candidate for CNS disorders involving mGlu_7_ dysfunction.

While the original characterization of CVN636 established its selectivity for mGlu_7/7_ over other mGlu receptor homodimers, it did not address whether CVN636 could also modulate mGlu_7_-containing heterodimers. This is a critical question, as it is now well established that mGlu receptors can form heterodimers in addition to homodimers, and that these heterodimeric complexes can exhibit pharmacological properties distinct from their homodimeric counterparts.^22–27^ mGlu_7_ has been shown to form heterodimers with mGlu_8_, mGlu_4_, mGlu_2_, and mGlu_3_ in heterologous cells,^28,29^ and evidence for the existence of some of these heterodimers in the brain is growing. For instance, mGlu_7/8_ heterodimers are supported by co-localization and functional data at hippocampal Schaffer collateral-CA1 synapses,^25,30,31^ while mGlu_7/4_ and mGlu_7/2_ heterodimers are plausible based on overlapping expression patterns and the demonstrated propensity of group II and III protomers to heterodimerize in native tissue.^32^ Notably the potency of glutamate at the mGlu_7_-containing heteromers is dramatically left shifted relative to the homomeric species.^25,27,33^ Given that CVN636 binds to the mGlu_7_ protomer and modulates mGlu_7/7_ homodimers, it is conceivable that it could also influence signaling at mGlu_7_-containing heterodimers, with important implications for interpreting its pharmacological effects both *in vitro* and *in vivo*.

To address these questions, we used two complementary bioluminescence resonance energy transfer (BRET)-based approaches: a cyclic adenosine monophosphate (cAMP) inhibition assay and our complemented donor-acceptor resonance energy transfer (CODA-RET) assay,^25,26^ the latter of which enables selective measurement of signaling from defined homo- or heterodimeric receptor complexes, to comprehensively characterize the pharmacology of CVN636 at mGlu_7/7_ homodimers and all major mGlu_7_-containing heterodimers. To our surprise, we found that CVN636 does not function as an allosteric agonist at mGlu_7/7_ homodimers, but instead acts as a PAM that enhances orthosteric agonist responses in an agonist-dependent manner. Furthermore, this PAM activity was selective for mGlu_7/7_ homodimers, with no detectable modulation at any mGlu_7_-containing heterodimer tested. These findings suggest that CVN636 is an mGlu_7/7_ homomer-selective PAM which has important implications for understanding both the mechanism of its *in vivo* efficacy and the therapeutic potential of targeting specific mGlu_7_ dimer populations.

## RESULTS

### CVN636 is a positive allosteric modulator of mGlu_7/7_ homodimers

To further characterize the pharmacology of CVN636 (**Figure 1A**) at mGlu_7_-containing receptors, we first sought to confirm that this compound acts as an allosteric agonist using a BRET-based cAMP inhibition assay (**Figure 1B**)^34^ that provides a downstream readout of mGlu_7_-mediated Gα_i/o_ activation. To our surprise, increasing concentrations of CVN636 alone, up to 100 µM, concentrations that are well in excess of the EC_50_ reported in Dickson et al., produced no detectable inhibition of cAMP at mGlu_7/7_ homodimers (**Figure 1C**, left). Moreover, comparison of the BRET signal at the maximal concentration tested revealed no difference between CVN636 and vehicle (**Figure 1C**, right), indicating that in our assay system CVN636 does not measurably activate mGlu_7/7_ homodimers on its own and therefore does not function as either an allosteric agonist or an ago-PAM.

**Figure 1.**
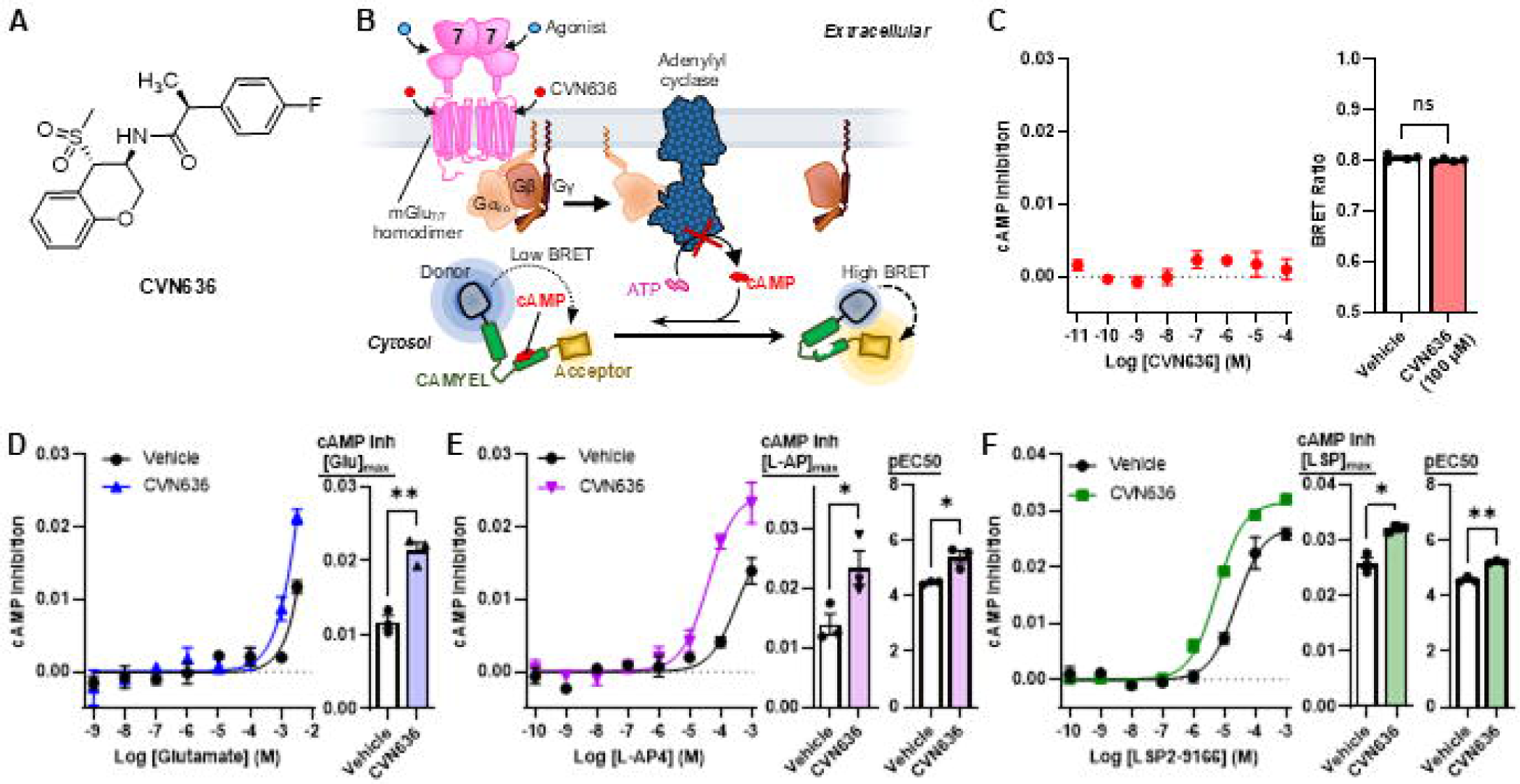
CVN636 does not activate mGlu_7/7_ homodimers alone but positively modulates orthosteric agonist responses. (**A**) Structure of CVN636. (**B**) Schematic of the BRET-based cAMP inhibition assay. Agonist-activated mGlu_7/7_ homodimers couple to and activate heterotrimeric Gα_i/o_ protein, leading to inhibition of adenylyl cyclase and decreased cAMP biosynthesis from adenosine triphosphate (ATP). The cAMP sensor using YFP-Epac-RLuc (CAMYEL) was used to detect changes in cAMP levels. (**C**) Left: CAMYEL measurements with mGlu_7/7_ homodimers showing a CVN636 concentration-response curve in the absence of orthosteric agonist. Right: comparison of the cAMP BRET signal at the maximal CVN636 concentration tested (100 μM) versus vehicle (DMSO). (**D–F**) CAMYEL measurements with mGlu_7/7_ homodimers showing concentration-response curves for (D) glutamate, (E) L-AP4, and (F) LSP2-9166 in the absence (DMSO vehicle) or presence of 50 µM CVN636 (left panels), with corresponding quantification of responses at maximal agonist concentration and/or pEC50 values shown in the right panels. For (C): n.s., p = 0.24. For (D): **p = 0.0022. For (E): cAMP inhibition at [L-AP4]_max_, *p = 0.047; pEC_50_, *p = 0.014. For (F): cAMP inhibition at [LSP2-9166]_max_, *p = 0.017; pEC_50_, **p = 0.0021. All comparisons by unpaired, two-tailed t-test. Symbols represent the mean drug-induced BRET response for three independent experiments performed in triplicate, and error bars represent ± SEM.

Given that CVN636 was efficacious in an *in vivo* model of alcohol use disorder,^21^ we next asked whether this compound might instead act as a PAM at mGlu_7/7_ by enhancing orthosteric agonist-mediated responses. We tested the effect of CVN636 on concentration-response curves for three orthosteric agonists: the endogenous agonist glutamate, and two selective synthetic group III mGlu receptor agonists, L-AP4 and LSP2-9166.^14,35,36^ For glutamate, consistent with the characteristically low affinity of mGlu_7_ for its endogenous agonist,^6,37^ detectable responses were observed only at high micromolar to millimolar concentrations, and the concentration-response curves did not reach saturation even at the highest concentration tested (**Figure 1D**, left). This precluded reliable estimation of potency or maximal response from curve fitting. Nevertheless, comparison of the cAMP inhibition signal at the highest glutamate concentration tested demonstrated a significant enhancement by CVN636 relative to vehicle (**Figure 1D**, right). For L-AP4 and LSP2-9166, CVN636 produced clear leftward shifts and increases in the maximal response of the concentration-response curves (**Figure 1E,F**, left panels), with quantification revealing significant increases in both potency and maximal response relative to vehicle (**Figure 1E,F**, right panels).

### CVN636 PAM activity requires orthosteric agonist binding

To confirm these findings using an independent and more proximal readout of receptor activation, we performed analogous experiments using our CODA-RET assay (**Figure 2A**), which can measure G protein recruitment to defined mGlu_7/7_ homodimers. Consistent with the cAMP results, CVN636 alone produced no detectable CODA-RET signal across the full concentration range tested (**Figure 2B**), with no difference from vehicle at the 100 μM concentration (**Figure 2B**, right), reinforcing the conclusion that CVN636 lacks intrinsic agonist activity in the CODA-RET assay as well. CVN636 also enhanced the activity of L-AP4 and LSP2-9166 in CODA-RET (**Figure 2D,E**), paralleling the effects observed in the cAMP assay. The glutamate concentration-response, however, was undetectable by CODA-RET (**Figure 2C**), likely because this assay measures G protein recruitment directly and lacks the signal amplification inherent in downstream cAMP readouts, rendering it insufficient to resolve the weak glutamate response at mGlu_7/7_ homodimers.

**Figure 2.**
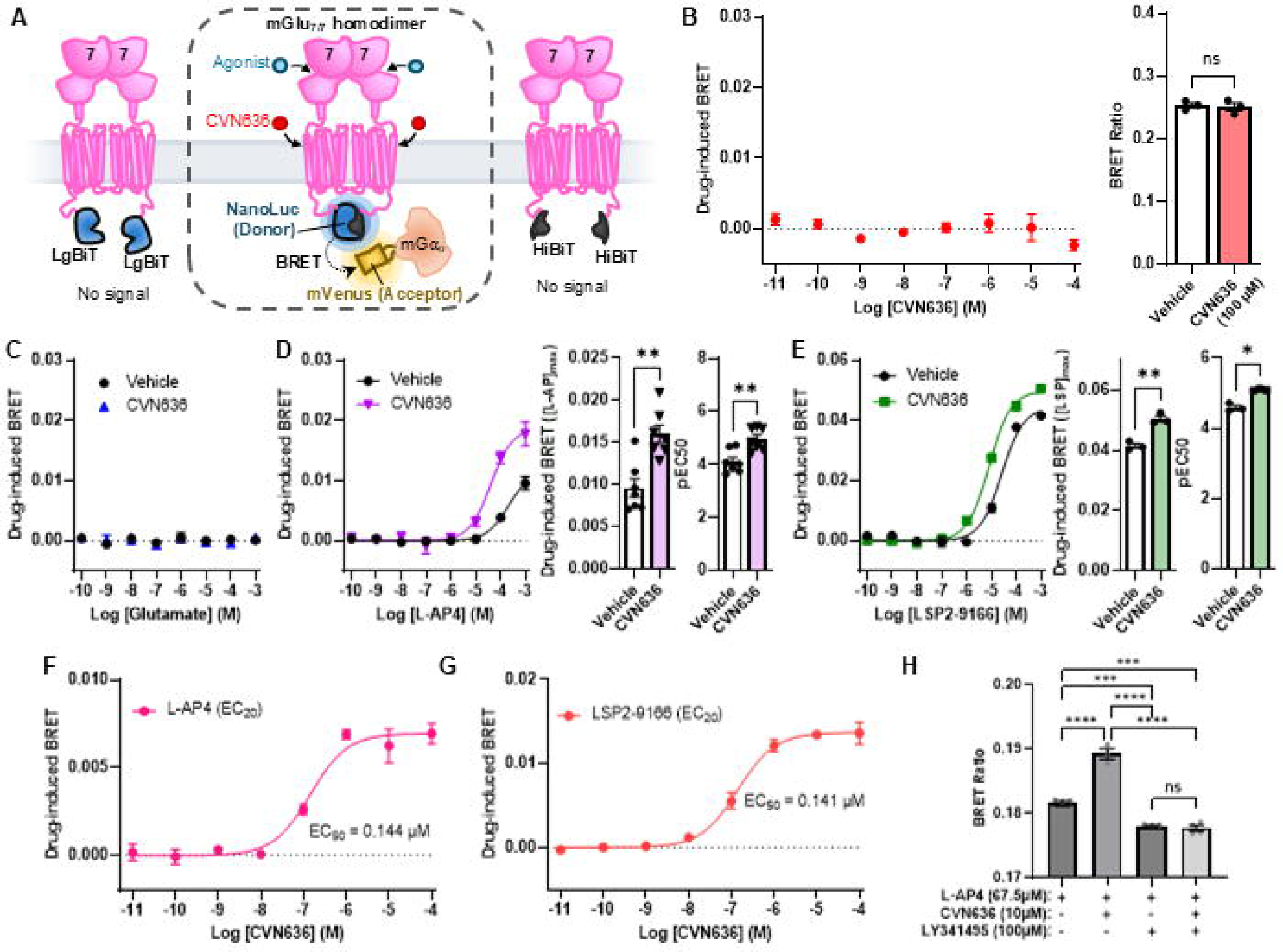
CVN636 potentiates orthosteric agonist-induced G protein recruitment at mGlu_7/7_ homodimers in an agonist-dependent manner. (**A**) Schematic of the CODA-RET assay performed with mGlu_7/7_ homodimers. mGlu_7_ protomers are fused to complementary LgBit and HiBit fragments of NanoLuc luciferase that produce bioluminescence only when a defined dimer assembles. BRET between the complemented NanoLuc (donor) and mVenus-tagged miniGα_o_ (acceptor) provides a readout of G protein recruitment to defined homodimers upon receptor activation. (**B**) Left: CODA-RET measurements showing a CVN636 concentration-response curve at mGlu_7/7_ homodimers in the absence of orthosteric agonist. Right: comparison of the CODA-RET signal at the maximal CVN636 concentration tested versus vehicle (DMSO). (**C–E**) CODA-RET measurements with mGlu_7/7_ homodimers showing concentration-response curves for (**C**) glutamate, (**D**) L-AP4, and (**E**) LSP2-9166 in the absence (DMSO vehicle) or presence of 50 µM CVN636 (left panels), with corresponding quantification of responses at maximal agonist concentration and/or pEC_50_ values shown in the right panels. (**F**) CVN636 concentration-response curve in the CODA-RET assay performed in the presence of L-AP4 near its EC20 (100 µM). (**G**) CVN636 concentration-response curve in the CODA-RET assay performed in the presence of a fixed EC_20_ concentration of LSP2-9166 (7.88 µM). (**H**) CODA-RET BRET ratios for mGlu_7/7_ homodimers stimulated with L-AP4 (67.5 µM) in the absence or presence of CVN636 (10 µM) and/or LY341495 (100 µM), as indicated. For (B): n.s., p = 0.75. For (D): CODA-RET at [L-AP4]max, **p = 0.0012; pEC□□, **p = 0.0028. For (E): CODA-RET at [LSP2-9166]max, **p = 0.0028; pEC□□, *p = 0.014. Comparisons in (B), (D), and (E) by unpaired, two-tailed t-test. For (H): ordinary one-way ANOVA with Tukey’s multiple comparisons test; L-AP4 vs. L-AP4 + CVN636, ****p = 1.8 × 10; L-AP4 vs. L-AP4 + LY341495, ***p = 8.7 × 10□□; L-AP4 vs. L-AP4 + CVN636 + LY341495, ***p = 5.9 × 10□□; L-AP4 + CVN636 vs. L-AP4 + LY341495, ****p = 3.4 × 10□□; L-AP4 + CVN636 vs. L-AP4 + CVN636 + LY341495, ****p = 2.9 × 10□□; L-AP4 + LY341495 vs. L-AP4 + CVN636 + LY341495, n.s., p = 0.99. Symbols represent the mean drug-induced BRET response for three independent experiments performed in triplicate, and error bars represent ± SEM.

Having established PAM activity in both assay platforms, we next sought to further characterize the pharmacological profile of CVN636. CVN636 concentration-response curves performed in the CODA-RET assay in the presence of fixed concentrations of L-AP4 or LSP2-9166 near their EC_20_ values revealed dose-dependent enhancement of the agonist-induced signal in both cases (**Figure 2F, G**), consistent with the expected behavior of a PAM whose activity depends on orthosteric agonist occupancy. The EC_50_ for CVN636 potentiation was similar for both agonists (140 nM). Finally, to determine whether CVN636’s potentiating effects require orthosteric agonist binding, we performed a competition experiment using the orthosteric antagonist LY341495 (**Figure 2H**).^14,38^ CVN636 enhanced the L-AP4-stimulated CODA-RET signal, but this enhancement was abolished when L-AP4 binding was competitively blocked by LY341495 (**Figure 2H**). This result is consistent with a PAM mechanism, as an allosteric agonist would be expected to activate the receptor independently of orthosteric site occupancy. Taken together, these data from two complementary assay platforms, measuring both downstream signaling and proximal G protein recruitment, indicate that CVN636 acts as a PAM, rather than an allosteric agonist, at mGlu_7/7_ homodimers.

### CVN636 does not modulate mGlu_7_-containing heterodimers

Having established that CVN636 acts as a PAM at mGlu_7/7_ homodimers, we next asked whether this compound could also modulate mGlu_7_-containing heterodimers. As mGlu_7_ has been reported to heterodimerize only with group II and III protomers, we first confirmed that CVN636 has no activity at the corresponding homodimers of these potential heterodimer partners. Using CODA-RET, we tested CVN636 at the other group II and III (except mGlu_6_) homodimers (**Supplemental Figure 1A–D**). CVN636 produced no detectable activation at these receptors (**Supplemental Figure 1E–H**), nor did it potentiate orthosteric agonist responses (**Supplemental Figure 1I–L**), confirming that CVN636 exhibits neither agonist nor PAM activity at these homomers. These results are consistent with the original characterization of CVN636, which also reported no activity at group II and III homodimers, nor at mGlu_6_ or the group I receptors mGlu_1/1_ and mGlu_5/5_.^21^

Using CODA-RET configured to selectively measure signaling from defined heterodimeric complexes, we tested CVN636 at mGlu_7/2_, mGlu_7/3_, mGlu_7/4_, and mGlu_7/8_ heterodimers (**Figure 3A–D**). Note that we did not test mGlu_7/6_, as mGlu_6_ expression is largely restricted to retinal ON bipolar cells and is not thought to be co-expressed with mGlu_7_ in brain circuits.^32,39^ CVN636 alone produced no detectable activation at any of these heterodimer combinations tested (**Figure 3E–H**), consistent with its lack of intrinsic agonist activity observed at mGlu_7/7_ homodimers. We then assessed whether CVN636 could potentiate orthosteric agonist responses at each heterodimer using L-AP4 and LSP2-9166. In contrast to the robust potentiation observed at mGlu_7/7_ homodimers (**Figure 2D, E**), CVN636 produced no detectable enhancement of either L-AP4 or LSP2-9166 concentration-response curves at mGlu_7/2_ (**Figure 3I, M**), mGlu_7/3_ (**Figure 3J, N**), mGlu_7/4_ (**Figure 3K, O**), or mGlu_7/8_ (**Figure 3L, P**) heterodimers. These results indicate that CVN636 PAM activity is selective for mGlu_7/7_ homodimers and does not extend to any of the mGlu_7_-containing heterodimers tested.

**Figure 3.**
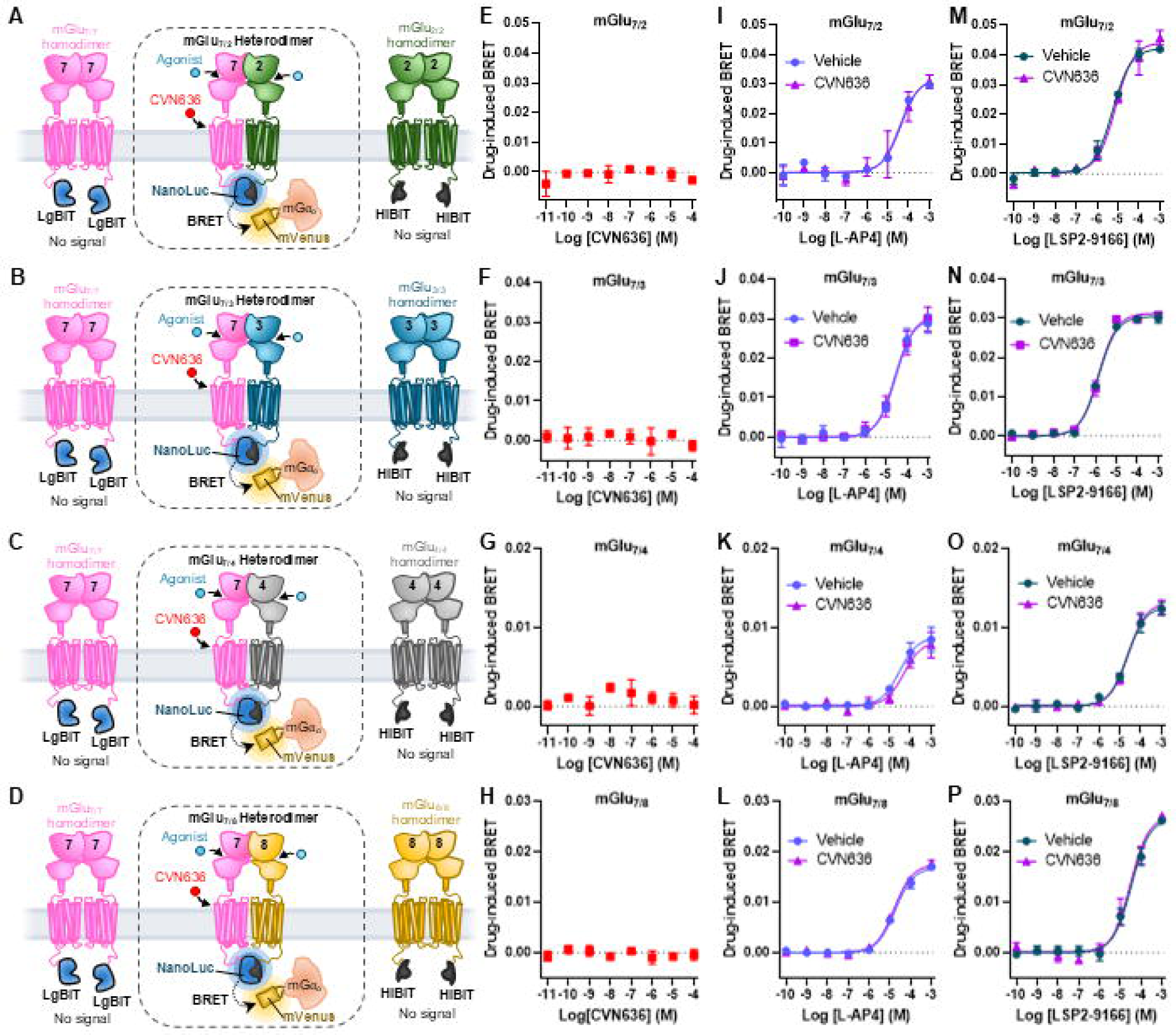
CVN636 does not modulate mGlu_7_-containing heterodimers. (**A–D**) Schematics of the CODA-RET assay configured to selectively measure signaling from defined (**A**) mGlu_7/2_, (**B**) mGlu_7/3_, (**C**) mGlu_7/4_, and (**D**) mGlu_7/8_ heterodimers. In each configuration, mGlu_7_ and partner protomers are fused to complementary fragments (LgBiT and HiBiT) of NanoLuc luciferase, such that only the defined heterodimer produces bioluminescence. Homodimers formed by protomers fused to non-complementing fragments (LgBiT/LgBiT or HiBiT/HiBiT) are optically silent and do not contribute to the BRET signal. (**E–H**) CODA-RET measurements showing CVN636 concentration-response curves at (**E**) mGlu_7/2_, (**F**) mGlu_7/3_, (**G**) mGlu_7/4_, and (**H**) mGlu_7/8_ heterodimers in the absence of orthosteric agonist. (**I–L**) CODA-RET measurements showing L-AP4 concentration-response curves at (I) mGlu_7/2_, (J) mGlu_7/3_, (K) mGlu_7/4_, and (L) mGlu_7/8_ heterodimers in the absence (DMSO vehicle) or presence of 50 µM CVN636. (**M–P**) CODA-RET measurements showing LSP2-9166 concentration-response curves at (**M**) mGlu_7/2_, (**N**) mGlu_7/3_, (**O**) mGlu_7/4_, and (**P**) mGlu_7/8_ heterodimers in the absence (DMSO vehicle) or presence of 50 µM CVN636. CVN636 showed no detectable agonist activity or potentiation of orthosteric agonist responses at any heterodimer tested. Symbols represent the mean drug-induced BRET response for three independent experiments performed in triplicate, and error bars represent ± SEM.

## DISCUSSION

In this study, we find that CVN636, previously reported as a potent allosteric agonist of mGlu_7_, acts as a positive allosteric modulator that requires orthosteric agonist co-activation and is selective for mGlu_7/7_ homodimers over all mGlu_7_-containing heterodimers tested. These findings have important implications for understanding mGlu_7_ pharmacology, the receptor’s role in brain circuits, and therapeutic strategies for targeting this receptor.

The original characterization of CVN636 concluded that it functions as an allosteric agonist based on its ability to inhibit cAMP production in CHO cells stably expressing mGlu_7_ in the absence of exogenously applied orthosteric agonist.^21^ In that study, receptor activation was measured as inhibition of forskolin-stimulated transcription from a CRE-luciferase reporter, read after a 5 h co-stimulation, a readout that both amplifies and integrates the signal. Consistent with substantial amplification, CVN636 was approximately 11-fold more potent in that assay than in the impedance-based measurement reported in the same study (7 nM versus 80 nM), the latter close to the 140 nM we measure by CODA-RET. Under those conditions, weak activity in the absence of added agonist could be read as agonism, whether it reflects very weak allosteric agonism, potentiation of constitutive activity, or potentiation of glutamate accumulating in the medium over the incubation.^40^ In our BRET-based cAMP and CODA-RET assays, where medium was exchanged immediately prior to compound addition and ambient glutamate was effectively controlled by excitatory amino acid transporter 3 (EAAT3) co-expression, CVN636 produced no detectable response in the absence of exogenously applied agonist, even at concentrations more than 7,000-fold above the reported EC_50_. However, CVN636 robustly potentiated responses to three distinct agonists: glutamate, L-AP4, and LSP2-9166, and this potentiation was abolished by the orthosteric antagonist LY341495, confirming an agonist-dependent PAM mechanism in the assays examined here. Dickson et al. also reported that LY341495 did not reduce the response to CVN636, which a competitive antagonist at twice its measured IC_50_ would not be expected to do against a signal integrated over such a long incubation.

The finding that CVN636 may act as a PAM rather than an allosteric agonist has important implications for understanding the *in vivo* efficacy of the compound. Dickson et al. demonstrated that CVN636 reduced ethanol self-administration in a rodent model of alcohol use disorder,^21^ an effect that, considering our findings, is likely to reflect potentiation of endogenous glutamate signaling at mGlu_7/7_ homodimers rather than direct receptor activation. This is a notable finding given the characteristically low affinity of mGlu_7_ for glutamate, which has led to the widely held view that this receptor functions as a synaptic ‘emergency brake,’ activated only under conditions of elevated glutamate release to limit sustained excitatory transmission.^6,8^ The low affinity has raised questions about whether PAMs targeting mGlu_7_-homomers could achieve meaningful potentiation of endogenous glutamate *in vivo*.^16^ However, peak glutamate concentrations in the synaptic cleft during transmission are estimated to reach 1–10 mM.^41,42^ These concentrations are within the range at which we observed CVN636-mediated potentiation of glutamate responses in our cAMP assay, suggesting that sufficient endogenous glutamate tone exists at mGlu_7_-expressing synapses to enable PAM activity. The *in vivo* efficacy of CVN636 therefore provides compelling evidence that mGlu_7/7_ homodimers are indeed engaged by endogenous glutamate under physiological or pathophysiological conditions and that their activity can be therapeutically modulated by positive allosteric modulation.

A striking finding of our study is that CVN636 PAM activity is selective for mGlu_7/7_ homodimers, with no detectable potentiation at mGlu_7/8_, mGlu_7/4_, mGlu_7/2_, or mGlu_7/3_ heterodimers. This degree of dimer-selective pharmacology, while perhaps surprising, is consistent with emerging evidence from our group and others that allosteric modulators can exhibit dramatically different activity profiles at mGlu homodimers versus heterodimers. We previously demonstrated that mGlu_4_ PAMs segregate into two classes based on their binding site within the transmembrane domain: those binding an upper allosteric pocket retain activity at mGlu_2/4_ heterodimers, whereas those binding a lower pocket are efficacious at mGlu_4/4_ homodimers but completely inactive at mGlu_2/4_ heterodimers.^26^ Similarly, we showed that mGlu_7_ NAMs exhibit differential activity at mGlu_7/7_ homodimers and mGlu_7/8_ heterodimers, with MMPIP blocking only homodimer responses while ADX71743 inhibits both.^25^ The complete loss of CVN636 activity at all four mGlu_7_-containing heterodimers tested here extends this principle and suggests that the allosteric binding site and/or the interprotomer communication pathway through which CVN636 exerts its PAM effects is fundamentally altered when the mGlu_7_ protomer is paired with a heterologous partner; the structural basis for this selectivity remains to be determined. It is possible that PAM activity requires the binding of two mGlu_7_ protomers.

The mGlu_7/7_ homomer selectivity of CVN636 positions it as a valuable pharmacological tool for dissecting the contributions of mGlu_7/7_ homodimers versus mGlu_7_-containing heterodimers in native brain circuits. At synapses where mGlu_7_ is co-expressed with other group II or III mGlu receptors, the receptor population likely comprises a mixture of homodimers and heterodimers, and traditional pharmacological approaches using orthosteric agonists or non-selective PAMs cannot distinguish between these species. CVN636, by virtue of its exclusive activity at mGlu_7/7_ homodimers, now provides a means to selectively probe homodimer function in electrophysiological and behavioral studies. When used in combination with heterodimer-active or heterodimer-selective compounds, CVN636 could enable pharmacological dissection of dimer-specific contributions to synaptic transmission and plasticity at circuits where multiple mGlu_7_-containing species are present. Such studies will be critical for determining whether homodimers and heterodimers play distinct or overlapping roles in the regulation of neurotransmission and for guiding the development of dimer-selective therapeutics.

In summary, our findings indicate that CVN636 is an mGlu_7/7_ homomer-selective PAM and suggest that positive allosteric modulation of mGlu_7/7_ homodimers can achieve therapeutic efficacy despite the receptor’s low glutamate affinity. These results highlight the importance of rigorously characterizing allosteric compounds across all physiologically relevant dimer combinations and open new avenues for precision pharmacology approaches that exploit dimer-specific drug action at mGlu receptors.

## EXPERIMENTAL PROCEDURES

### Compounds

L-AP4 and LY341495 were purchased from Tocris. LSP2-9166 was synthesized at the Vanderbilt Molecular Design and Synthesis Core using methods reported in Acher et al.^43^ CVN636 was a gift from Cerevance and further purchased from MedChemExpress.

### Construction and transfection of expression vectors

cDNA for rat mGlu_2_, mGlu_3_, mGlu_4_, mGlu_7_ and mGlu_8_ were N-terminally tagged with a hemagglutinin epitope tag using standard molecular biology procedures. cDNAs encoding the split fragments of Nano luciferase (Nanoluc, Promega), LgBit (residues 1-158) or HiBit (residues 159-169: VSGWRLFKKIS), were fused in frame to the C terminus of related mGlu receptor sequences following the linker “EFPPARATLPPARATLE” in the pcDNA3.1 vector. The following G protein constructs in pcDNA3.1 were also used: A truncated Gαo subunit fused to mVenus, mVenus-miniGo, as well as untagged G_αi_, G_β1_, and G_γ2_. The cDNA encoding the cAMP sensor using <u>Y</u>FP-Epac-R<u>L</u>uc (CAMYEL) and Excitatory amino acid transporter 3 (EAAT3) were described previously.^34,44^ The integrity of all the constructs was confirmed with sequencing analysis.

### Cell culture and transfections

All cell-based assays were carried out in 293T cells (ATCC, CRL-3216) from embryonic kidney tissue of Homo sapiens (HEK293T). The cells are female in origin and were authenticated by the manufacturer using short tandem repeat analysis to determine species and unique DNA profile. The cells were maintained in Dulbecco’s modified Eagle’s medium (DMEM), high glucose (Gibco) supplemented with 10% fetal bovine serum (Corning Inc.) and 1% penicillin-streptomycin (Corning Inc.) at 37 °C with 5% CO_2_. Prior to transfection, HEK293T grew to ∼70% confluency in 10 cm tissue culture dishes. The HEK293T cells used in this study tested negative for mycoplasma contamination.

For the CODA-RET assays, the pcDNA 3.1 (+) plasmids encoding mGlu_2_, mGlu_3_, mGlu_4_, mGlu_7_ or mGlu_8_ protomers (mGlu_2_ homodimer expression: 250ng mGlu_2_-LgBit and 250ng mGlu_2_-HiBit; mGlu_3_ homodimer expression: 50ng mGlu_3_-LgBit and 50ng mGlu_3_-HiBit; mGlu_4_ homodimer expression: 250ng mGlu_4_-LgBit and 250ng mGlu_4_-HiBit; mGlu_7_ homodimer expression: 100ng mGlu_7_-LgBit and 100ng mGlu_7_-HiBit; mGlu_8_ homodimer expression: 50ng mGlu_8_-LgBit and 50ng mGlu_8_-HiBit; mGlu_7/2_ heterodimer expression: 200ng mGlu_7_-LgBit and 250ng mGlu_2_-HiBit; mGlu_7/3_ heterodimer expression: 200ng mGlu_7_-LgBit and 50ng mGlu_3_-HiBit; mGlu_7/4_ heterodimer expression: 200ng mGlu_7_-LgBit and 250ng mGlu_4_-HiBit; mGlu_7/8_ heterodimer expression: 200ng mGlu_7_-LgBit and 100ng mGlu_8_-HiBit;) were co-transfected with 5µg miniG_αo_ tagged with mVenus and 1µg EAAT3 into HEK293T cells using PEI Max 4000 at a ratio of 9µg PEI Max per 1µg total plasmid. Transfected cells were incubated at 37°C with 5% CO_2_ for 48h. Note that prior to experiments, the cells were incubated with DMEM, high glucose, GlutaMAX supplement, pyruvate (Gibco) in the absence of fetal bovine serum for 1h at 37°C with 5% CO_2_.

For the cAMP assay, the pcDNA 3.1 (+) plasmid encoding mGlu_7_-LgBiT (mGlu_7_ homodimer expression: 4µg mGlu_7_-LgBiT), which cannot complement to produce functional Nanoluc, was co-transfected with 6 µg of the CAMYEL, 1 µg G_αi_, G_β1_, G_γ2_ and EAAT3 plasmid into HEK293T cells using PEI Max 4000 at a ratio of 2µg PEI Max per 1µg total plasmid. The same expression protocol described for the CODA-RET assay was followed for the cAMP assay.

### CODA-RET assay

After expression, the cells were dissociated from the 10 cm plate using enzyme free dissociation solution (Millipore-Sigma). After dissociation, the cells were collected and spun at 1000g for 5min. The supernatant was discarded, and the cells were washed twice with pre-warmed Dulbecco’s phosphate-buffered saline (DPBS). Approximately 150,000 cells per well were distributed in black frame, white well 96-well plates (PerkinElmer). 5 µM furimazine, the substrate for Nanoluc, was added to each well and immediately after the addition of furimazine, pre-warmed DPBS containing the indicated mGlu receptor agonist and PAM or vehicle was loaded to each well. After five minutes of ligand stimulation at 37°C, the fluorescence and luminescence signals were quantified using a Pherastar FS plate reader (BMG Labtech). The BRET ratio was determined by calculating the ratio of mVenus signal (535 nm) divided by the Nanoluc (485 nm) signal. The results are expressed as the BRET change produced by the corresponding ligands.

### cAMP assay

The cells were dissociated, washed, and plated as described above for the CODA-RET assay. Approximately 60,000 cells per well were distributed in 96-well plates and stimulated by indicated ligands dissolved in pre-warmed DPBS containing 10μM forskolin for 30min at 37°C. 5 µM Coelenterazine H was added to each well 5 min before reading. The fluorescence and luminescence were quantified and data analyzed as described above.

### Plotting and Statistics

Data from the CODA-RET and other BRET-based live cell assays were plotted and fit using GraphPad Prism (GraphPad Software). The dose–response curves of agonist and PAM responses were fitted by nonlinear regression to the log(agonist) vs. response model with a standard Hill slope equal to 1 and the bottom fit constrained to 0. EC_20_ concentrations were calculated from the concentration-response curves measured here; L-AP4 was used at 100 µM, near its calculated EC_20_ of 70 µM.

This article contains supporting information.

## DATA AVAILABILITY

All data are contained within the manuscript and are available upon request from Wesley B. Asher and Jonathan A. Javitch

## ACKNOWLEDGEMENTS

We thank Jiya Singh for technical assistance.

## FUNDING AND ADDITIONAL INFORMATION

This work was supported by National Institutes of Health grants R01-NS132060 (J.A.J.), the St. Jude Children’s Research Hospital GPCR Collaborative (J.A.J.), the Hope for Depression Research Foundation (J.A.J.), and Miriam’s Magical Memorial Mission (J.A.J.).

## ABBREVIATIONS

BRET: bioluminescence resonance energy transfer;
cAMP: cyclic adenosine monophosphate;
CODA-RET: complemented donor-acceptor resonance energy transfer;
CVN636: (αS)-N-[(3S,4S)-3,4-Dihydro-4-(methylsulfonyl)-2H-1-benzopyran-3-yl]-4-fluoro-α-methyl benzeneacetamide;
DMSO: dimethyl sulfoxide;
EC_50_: half maximal effective concentration;
Emax: maximal effect;
Gα_i/o_: inhibitory G protein alpha subunit;
L-AP4: L-2-amino-4-phosphonobutyric acid;
LSP2-9166: (2S)-2-amino-4-[[[4-(carboxymethoxy)-3-(trifluoromethoxy) phenyl]-hydroxymethyl]-hydroxy phosphoryl]butanoic acid;
mGlu: metabotropic glutamate;
PAM: positive allosteric modulator;
pEC_50_: negative logarithm of the half maximal effective concentration

