## Supplemental Figure 1 for "CVN636 potentiates orthosteric agonist responses selectively at mGlu_7/7_ homodimers"

- 1.) Department of Psychiatry, Columbia University Vagelos College of Physicians and Surgeons, New York, NY 10032, USA
- 2.) Division of Preclinical Research, New York State Psychiatric Institute, New York, NY 10032, USA
- 3.) Department of Pharmacology, Warren Center for Neuroscience Drug Discovery, Vanderbilt Brain Institute, Vanderbilt Institute of Chemical Biology, Vanderbilt Institute for Translational Advances, Vanderbilt University, Nashville, TN 37232, USA
- 4.) Vanderbilt Kennedy Center, Vanderbilt University Medical Center, Nashville, TN 37232, USA
- 5.) Department of Pharmacology, Columbia University Vagelos College of Physicians and Surgeons, New York, NY 10032, USA

### Contents

Supplemental Figure 1. CVN636 does not modulate group II and III mGlu homodimers.

### SUPPLEMENTAL FIGURE

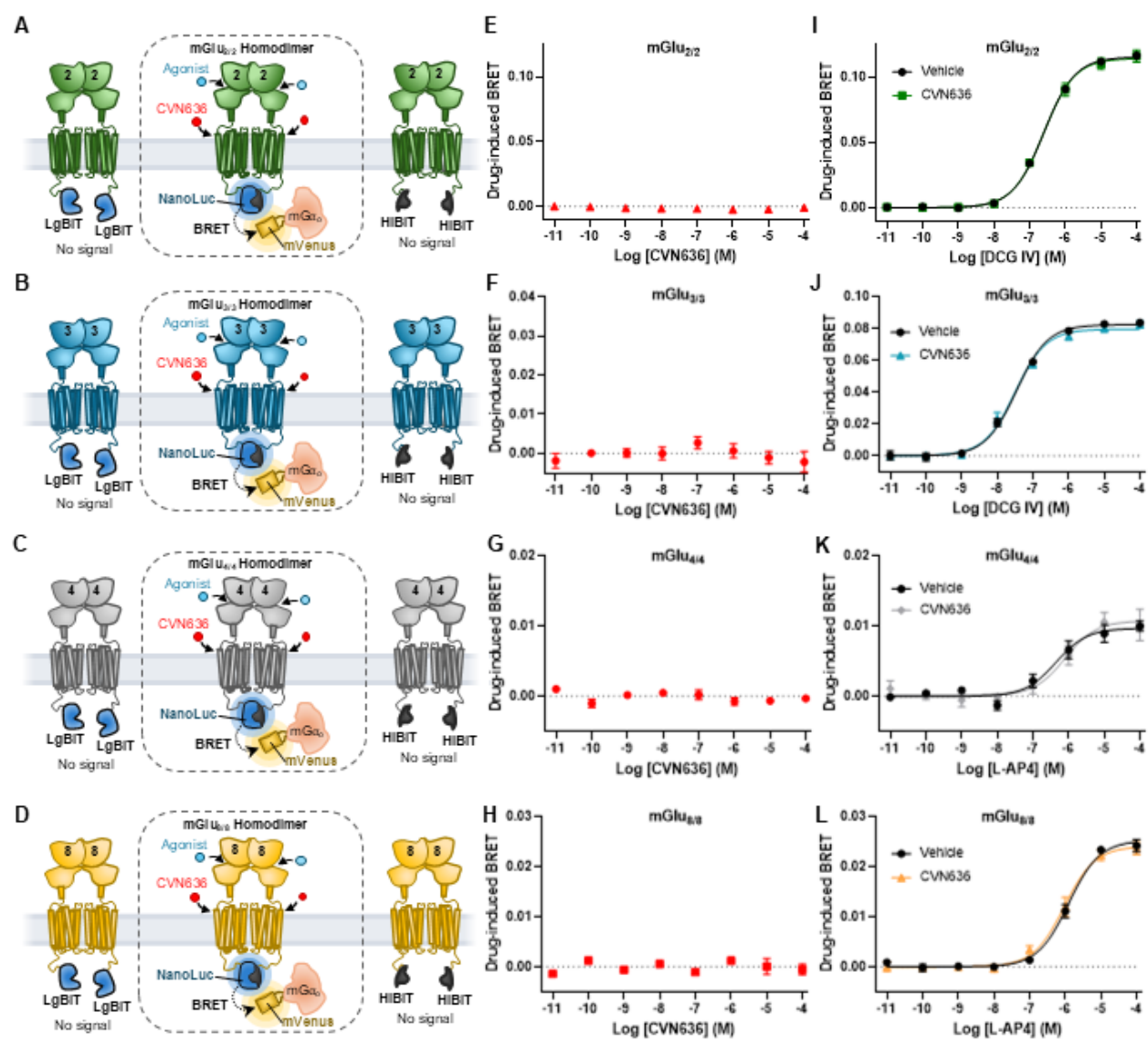

**Supplemental Figure 1. CVN636 does not modulate group II and III mGlu homodimers.**

(A–D) Schematics of the CODA-RET assay configured to measure signaling from defined (A) mGlu<sub>2/2</sub>, (B) mGlu<sub>3/3</sub>, (C) mGlu<sub>4/4</sub>, and (D) mGlu<sub>8/8</sub> homodimers. (E–H) CODA-RET measurements showing CVN636 concentration-response curves at (E) mGlu<sub>2/2</sub>, (F) mGlu<sub>3/3</sub>, (G) mGlu<sub>4/4</sub>, and (H) mGlu<sub>8/8</sub> homodimers in the absence of orthosteric agonist. (I–L) CODA-RET measurements showing orthosteric agonist concentration-response curves at (I) mGlu<sub>2/2</sub> (DCG-IV), (J) mGlu<sub>3/3</sub> (DCG-IV), (K) mGlu<sub>4/4</sub> (L-AP4), and (L) mGlu<sub>8/8</sub> (L-AP4) homodimers in the absence (vehicle) or presence of 50 μM CVN636. CVN636 showed no detectable agonist activity or potentiation of orthosteric agonist responses at any homodimer tested. Symbols represent the mean drug-induced BRET response for three independent experiments performed in triplicate, and error bars represent ± SEM.
